# Adaptive strategies of the candidate probiotic *E. coli* Nissle in the mammalian gut

**DOI:** 10.1101/364505

**Authors:** Nathan Crook, Aura Ferreiro, Andrew J. Gasparrini, Mitchell Pesesky, Molly K. Gibson, Bin Wang, Xiaoqing Sun, Zevin Condiotte, Stephen Dobrowolski, Daniel Peterson, Gautam Dantas

## Abstract

Probiotics are living microorganisms that are increasingly used as gastrointestinal therapeutics by virtue of their innate or engineered genetic function. Unlike abiotic therapeutics, probiotics can replicate in their intended site, subjecting their genomes and therapeutic properties to natural selection. By exposing the candidate probiotic *E. coli* Nissle (EcN) to the mouse gastrointestinal tract over several weeks, we uncovered the consequences of gut transit, inter-species competition, antibiotic pressure, and engineered genetic function on the processes under selective pressure during both within-genome and horizontal evolutionary modes. We then show the utility of EcN as a chassis for engineered function by achieving the highest reported reduction in serum phenylalanine levels in a mouse model of phenylketonuria using an engineered probiotic. Collectively, we demonstrate a generalizable pipeline which can be applied to other probiotic strains to better understand their safety and engineering potential.

## Introduction

The human gut microbiota is the collection of microbes which have evolved to live in the human gastrointestinal tract (Huttenhower et al., 2012). The microbiota’s metagenome complements the human genome by assisting in, among other functions, metabolism of dietary fiber (El Kaoutari et al., 2013), synthesis of vitamins (Backhed et al., 2005), and exclusion of pathogens (Theriot et al., 2014). Chemical and/or microbial perturbations to this community can result in dysbiotic community compositions, defined by their negative impact on human health (Sampson et al., 2016; Smith et al., 2013; Tamboli et al., 2004; Turnbaugh et al., 2006). Correction of specific dysbioses through administration of probiotics has shown therapeutic promise (Sartor, 2005). Probiotics are live microorganisms that, when consumed in sufficient quantities, provide health benefits to the host, including enhancement of epithelial barrier function, blocking of pathogen binding, and vitamin synthesis (Hill et al., 2014). To improve efficacy and expand therapeutic scope, probiotics can be genetically engineered, with recent work demonstrating preclinical efficacy against infectious and metabolic diseases (Durrer et al., 2017; Hwang et al., 2017; Mansour and Abdelaziz, 2016; Palmer et al., 2018). Unlike conventional small-molecule (abiotic) therapeutics, wild type or engineered probiotics replicate in the gut and are therefore subject to genomic mutation and natural selection, potentially to the detriment of their intended therapeutic effect and safety profile. Indeed, both commensal and pathogenic microbes have been shown to adapt in a host-specific manner during gut passage (Lieberman et al., 2014; Zhao et al., 2018). Clinical use of probiotics therefore demands a thorough understanding of their *in vivo* evolutionary trajectories. In this study, we focus on the *in vivo* evolutionary trajectories of the candidate probiotic *Escherichia coli* Nissle 1917 (EcN), an *E. coli* clade B2 probiotic [Figure S2]. EcN has a long history of probiotic usage without evolution of pathogenic phenotypes (Wassenaar et al., 2015; Westendorf et al., 2005), has demonstrated efficacy against inflammatory bowel disease (Scaldaferri et al., 2016), and has been gaining increased attention as a “chassis” for engineered therapeutic function (Hwang et al., 2017; Palmer et al., 2018).

One example of a disorder which could be treated with engineered probiotics is phenylalanine-hydroxylase-deficient phenylketonuria (PKU), an inborn error of host metabolism. Individuals with phenylketonuria lack some or all function of the phenylalanine hydroxylase enzyme (PAH), leading to accumulation of neurotoxic phenylalanine (Phe) levels which, if left untreated, impair neurocognitive development and lead to severe intellectual disability (Nardecchia et al., 2015; Regier and Greene, 2000). The primary therapy for PKU is dietary restriction through use of low Phe and low protein medical foods to achieve serum Phe concentrations in the range of 120-360 µmol/L (Vockley et al., 2014). However, the efficacy of dietary restriction is substantially hindered by poor compliance (MacDonald et al., 2010; MacLeod and Ney, 2010). The lone FDA approved therapeutic alternative is sapropterin, but this is effective only for a subset of PKU patients (Vockley et al., 2014). While an enzyme replacement therapy (Pegvaliase) (Eavri and Lorberboum-Galski, 2007) has undergone promising Phase III clinical trials (Charlesworth, 2016), this option is expensive and requires continual administration. Recent work establishing efficacy of an engineered *Lactobacillus* strain in a murine model of PKU demonstrates that engineered probiotics could innovatively address shortcomings in current therapies (Durrer et al., 2017). However, the level of reduction of murine serum Phe concentrations achieved in this *Lactobacillus* study was substantially lower than those reported with the injected enzyme therapy (Sarkissian et al., 2008), motivating further optimization of engineered probiotics for PKU. We hypothesized that EcN, with established genetic tools enabling high-output enzyme production, would provide an alternative and potentially superior host for therapeutic degradation of gut Phe through regulated expression of a heterologous phenylalanine-ammonia lyase (PAL). PAL converts Phe into ammonia and trans-cinnamic acid, both of which are nontoxic and readily excreted (Sarkissian and Gamez, 2005). Additionally, we hypothesized that, due to their differing serum Phe levels, the optimal probiotic PAL expression level would differ between male and female PKU mice.

Prior work has uncovered the unique *in vivo* evolutionary trajectories (relative to *in vitro*) of commensal, laboratory, and pathogenic strains of *E. coli* in the mammalian gut under specific dietary conditions and gut microbiome architectures (Barroso-Batista et al., 2014; Ghalayini et al., 2018; Gumpert et al., 2017; Karami et al., 2007; Lescat et al., 2017; Lourenco et al., 2016). Nutrient availability and inter-species metabolic dependencies are major selective pressures for the gut microbiome (Hoek and Merks, 2017; Rakoff-Nahoum et al., 2016; Zhao et al., 2018). A study of a large human cohort (N > 1000) demonstrated that host diet dominates host genetics as a driver of the composition and function of the gut microbiota (Rothschild et al., 2018). It is also well-known that engineered metabolism can dramatically compromise microbial growth (Fletcher et al., 2016; Lechner et al., 2016), and in general may alter the primary selective pressures relative to wild type. We therefore set out to investigate the effects of host diet, microbiome complexity, and engineered metabolism on the *in vivo* evolutionary trajectories of wild type and engineered EcN, using mice as a model of the mammalian gut (Reyes et al., 2013; Ridaura et al., 2013) and heterologous expression of PAL as a model of engineered probiotic metabolism.

Probiotics face a variety of selective pressures in the mammalian gut that may lead to *in vivo* evolution. First, EcN cannot natively degrade many of the complex polysaccharides present in the intestine (Fabich et al., 2008; Hoskins et al., 1985), and is dependent on other species for production of metabolizable di- and mono-saccharides (Conway and Cohen, 2015), which may lead to adaptations in carbohydrate utilization. Second, the high prevalence of antagonistic interspecies interactions in the gut (Wexler et al., 2016) may drive adaptation in EcN’s encoded colicins (Patzer et al., 2003). Third, since EcN normally resides in sessile multi-species biofilms on the outer mucin layer (Adediran et al., 2014; Conway and Cohen, 2015), introduction to the murine gut may drive adaptation in adhesins, increasing their specificity to mouse mucins. Finally, expression of a Phe-degrading enzyme may impose a metabolic burden. For these reasons, we hypothesized that the primary genomic adaptations (mutations) which EcN would undergo in the mouse gut could involve i) carbohydrate utilization, ii) inhibitory interactions with other microbes, iii) motility/adhesion, or iv) inactivation of engineered function.

For gut commensals, genetic adaptation can occur through genomic SNPs or indels, as well as horizontal gene transfer (evolutionary scales 1 and 2, respectively, as described in (Ferreiro et al., 2018)). Horizontal gene transfer is a major mechanism driving bacterial evolution (Ferreiro et al., 2018; Rodriguez-Beltran et al., 2018; Soucy et al., 2015), occurring more rarely than single nucleotide polymorphisms (SNPs) and insert/deletion events but accounting for greater net genetic distance on timescales shorter than bacterial speciation, and frequently associated with key functional determinants such as antibiotic resistance (Smillie et al., 2011). Bacterial strains differ in their capacity to engage in HGT, due to origin of replication incompatibilities and lack of natural competence (Thomas and Nielsen, 2005). In this study, we describe the fitness and adaptation of EcN in the murine gut using both whole-genome sequencing of *in vivo* adapted isolates (to recover changes in the EcN genome) and high-throughput functional metagenomic selections (to simulate horizontal gene transfer). In each case, replicate mice were fed diverse diets including a standard mouse diet, a high-fat Western diet [Tables S1-S2], and diets supplemented with prebiotics. Further, we included mice with a spectrum of background microbiota complexities, ranging from germ-free mice mono-colonized with EcN, germ-free mice colonized with synthetic biota of intermediate complexity, and conventional mice with antibiotic-perturbed or unperturbed complex microbiota [Figure S1]. Using this reductionist approach, we elucidated the impact of host diet and microbial competition on the *in vivo* adaptation of EcN. In total we analyzed 225 longitudinal samples from functional metagenomic selections in 33 mice [Table 1], and 401 EcN isolates from 20 mice [Table S3], comprising the most comprehensive study of *in vivo* probiotic evolution to date. Finally, using engineered probiotic EcN, we achieve the highest reported reduction in murine serum Phe levels from an engineered probiotic, and describe genomic changes occurring *in vivo* during course of treatment.

**Table 1.** **Summary of Functional Metagenomic Selection Experiments.** See also Table S1, Table S2, and Table S4 for descriptions of the diet compositions and the 13-member community.

| Microbiome | Diet | # Mice |
| --- | --- | --- |
| Germ-free | Mouse | 6 |
| Germ-free | Human | 5 |
| Germ-free | Human+Dextrin | 5 |
| Germ-free | Human+Inulin | 5 |
| Germ-free | Human | 3 |
| 13-member | Human | 3 |
| 13-member | Human+Dextrin | 3 |
| 13-member | Human+Inulin | 3 |

## Results

### Interrogation of HGT-mobilizable functions

To identify functions that may be acquired for improved EcN fitness in the mammalian gut, we carried out functional metagenomic selections where metagenomic insert libraries heterologously expressed in EcN were passaged through mouse gastrointestinal tracts for up to five weeks (Gibson et al., 2014). By creating a functional metagenomic library, we were able to bypass the normal limitations and timescale of HGT and ask: if EcN could receive any genetically encoded function from any member of a healthy gut microbiome independent of mode of HGT, what functions would provide the greatest fitness advantage? By competing millions of recombinant EcN strains against each other *in vivo*, we expected to observe significant enrichment of a subpopulation of strains with DNA inserts conferring EcN a fitness advantage. These functional metagenomic selections served as an upper bound proxy for the potential impact of horizontal gene transfer on the fitness of EcN in the gut. The source of our metagenomic DNA was stool from two healthy human infants and six healthy human mothers, to approximate the diversity of microbial taxa present in the gut through different life stages (Moore et al., 2015). Approximately 10^6^ unique 2-5kb fragments of metagenomic DNA were cloned into EcN, generating a library >10^9^ bp in size, which is equivalent to hundreds of bacterial genomes.

To more easily parse the selective pressures placed on this functional metagenomic library, we performed *in vivo* selections using a reductionist approach. That is, we began with the *in vivo* setting with the fewest selective factors: the germ-free gut of mice fed only standard mouse chow and water. Subsequently, we carried out functional selections in mice pre-colonized with a defined community of 13 representative human gut bacterial species (Goodman et al., 2011), conventional mice that had had their microbiota depleted through streptomycin treatment, and untreated conventional mice. We also carried out selections in mice fed either standard mouse chow, a high-fat Western diet, or a high-fat diet supplemented with either dextrin or inulin, two prebiotic polysaccharides. We specifically hypothesized that in EcN library mono-colonized mice, we would select for enzymes capable for degrading complex polysaccharides, a function that *E. coli* typically depends on gut anaerobes to carry out. Conversely, we hypothesized that co-colonization with a defined bacterial community would relieve this selective pressure, which would be reflected in a greater diversity of enriched DNA inserts.

### Selection for carbohydrate utilization in the germ-free gut

Metagenomic libraries were delivered to six germ-free C57BL/6 mice via oral gavage. These mice consumed a standard mouse diet (MD) composed mainly of plant-based carbohydrates and protein [Tables S1-S2]. Over 5 weeks of colonization, we observed a striking reduction in the diversity of metagenomic inserts in mouse feces. In all 6 mice, the initial ∼10^6^ metagenomic fragments were reduced to the same 5 metagenomic inserts. [Figure 1b]. Supporting our hypothesis that in the absence of microbial taxa capable of degrading complex polysaccharides, the main selective pressure on EcN in the gut is polysaccharide degradation, four out of the five inserts contained a predicted glycosyl hydrolase family 32 (GH32) enzyme. GH32 is a large protein family involved in the hydrolysis of glycosidic bonds within polysaccharides. Several of these inserts also contained predicted transporters and other enzymes involved in central carbon metabolism [Figure 1g]. To experimentally validate our hypothesis that these *in vivo*-enriched inserts directly enhance the metabolic activity of EcN, we cloned these metagenomic fragments into wild type EcN and tested additional carbon source utilization conferred by these fragments using *in vitro* phenotypic microarrays [Supplementary File 1].

**Figure 1.**
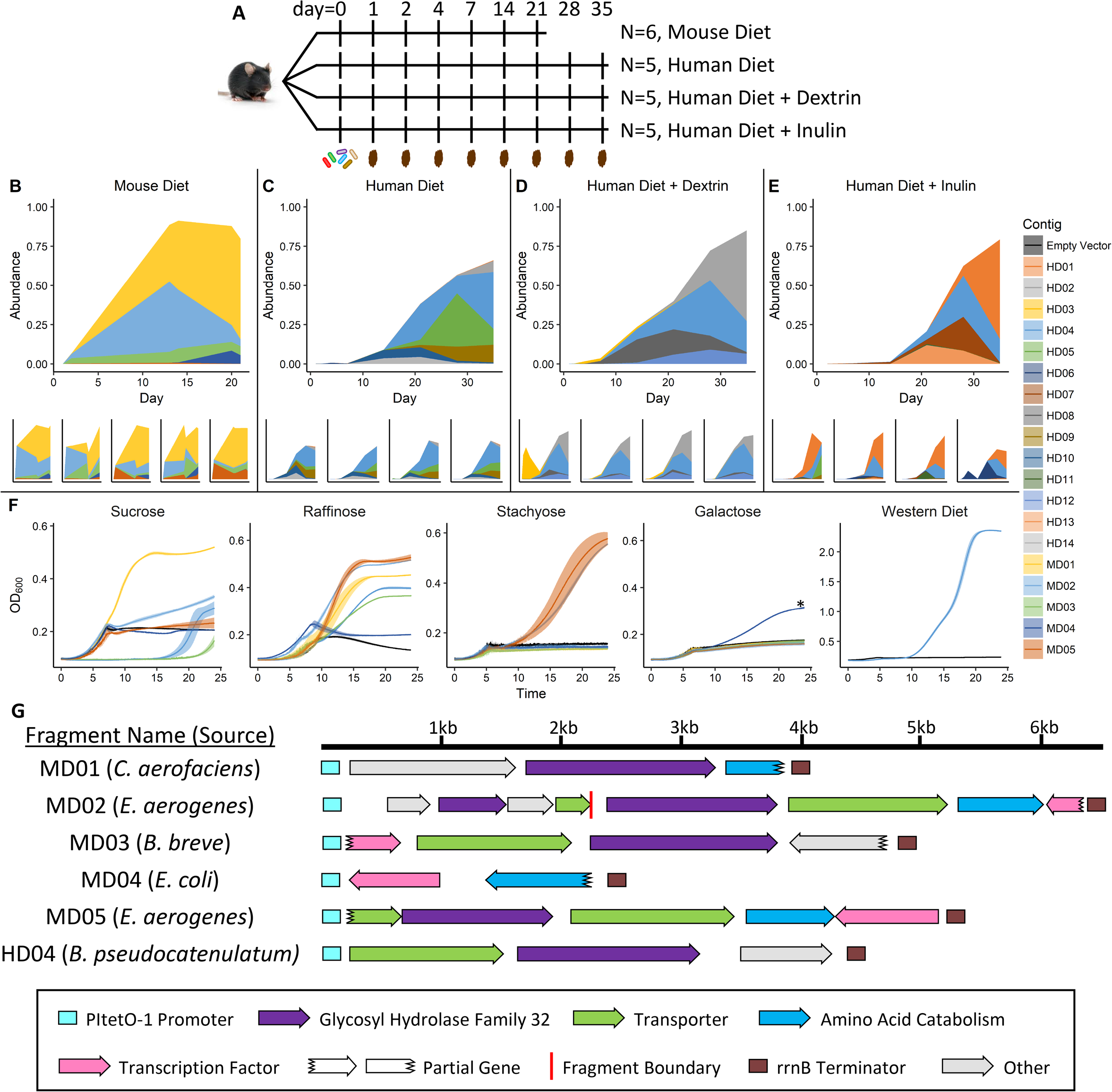
Functional metagenomic selections in EcN mono-colonized mice. **A)** Experimental design and sampling timeline. Mice were gavaged on day 0. N: number of replicate mice. **B-E)** Relative abundance of metagenomically sourced fragments in the EcN population in replicate mice fed standard mouse diet and water (B), a high-fat human diet and water (C), a high-fat human diet and dextrin (D), or a high-fat human diet and inulin (E).**F)** Growth curves of select metagenomic fragments enriched in the functional selections depicted in B-E on minimal media supplemented with different carbon sources. * P < 6×10-3, Student’s t-test comparing time taken to reach OD_600_=0.25 between WT and MD04. **G)** Annotations of the metagenomic fragments enriched in the functional selection depicted in B, and the generalist fragment enriched in the functional selections depicted in C-E. MD: fragments enriched in mice fed a Mouse Diet; HD: fragments enriched in mice fed a Human Diet. See also Figures S1-S3, and Table S6.

GH32 is reported to be involved in sucrose utilization. Commensal *E. coli*, including EcN, are unable to consume sucrose (Conway and Cohen, 2015), and each GH32-containing fragment enabled growth on this substrate [Figure 1f]. Similarly, several GH32-containing fragments enabled growth on longer-chain oligosaccharides containing sucrose as a terminal moiety, including raffinose and stachyose [Figure 1f]. The growth rate of EcN strains containing 2 different GH32-containing fragments (MD02 and MD05) was higher on these longer-chain oligosaccharides than sucrose, indicating likely specialization of these enzymes toward these molecules. The one fragment we isolated from this experiment which did not contain GH32 (fragment MD04) instead contained a transcription factor which was identical (100% nucleotide identity) to the transcriptional regulator gadX in *E. coli* Nissle. This protein plays a major role in regulating acid tolerance by regulating the expression of the *gad* operon, which consumes intracellular H+ during glutamate metabolism (Tramonti et al., 2002; Tucker et al., 2003). It has been previously shown that cytosolic pH regulates glycolytic flux as mediated by vacuolar ATPases - proton pumps that sense intracellular pH and cease activation of the cAMP-dependent protein kinase A under acidic conditions (Dechant et al., 2010). Supporting this hypothesized role, *in vitro* recombinant expression of this fragment in EcN significantly reduced lag phase when switching from LB to galactose-containing minimal media [Figure 1f].

Our observations regarding the similarity of recovered fragment functions were confirmed after assigning Gene Ontology (GO) annotations to these 5 fragments. In agreement with our observations, >87% of functionally selected fragments were annotated as belonging to the GO pathway “Carbohydrate Metabolic Process.” The next most abundant GO pathways were “Phosphorylation” followed by “Carbohydrate Transmembrane Transport” and “Transcription, DNA-templated”. Together, these results indicated that the dominant selective pressure on EcN in the germ-free mouse gut is for enzymatic degradation and utilization of long-chain polysaccharides.

### Diet-dependent selection

We hypothesized that because selective pressure for long-chain polysaccharide degradation was so strong in the EcN mono-colonized mouse gut, we would observe polysaccharide-specific enrichment of DNA inserts in response to the presence to prebiotics. To test this hypothesis, we gavaged EcN containing the same metagenomic library to 3 groups of 5 germ-free C57BL/6 mice. All 15 mice were maintained on a high-fat, high-carbohydrate diet which mimics the diet of Western humans (HD) (Plump et al., 1992). One group of mice was maintained on the Western diet alone, while the other two were supplemented with the prebiotics inulin or dextrin in the drinking water. Inulin is a fructose polysaccharide with chain-terminating glucosyl moieties and is not degradable by wild type EcN, while dextrin is a glucose polysaccharide that can support limited EcN growth, e.g. 20-30% of the cell density that glucose can support under aerobic conditions (Fabich et al., 2008), [Figure S3]. Over 5 weeks, we again observed strong convergence toward ≤6 metagenomic fragments in each dietary condition [Figure 1c-1e]. Supporting our diet-dependency hypothesis, the collection of enriched fragments in each condition was different. However, one “generalist” metagenomic fragment was enriched in all three groups of mice (fragment HD04). This fragment was unique among the enriched fragments in that it contained a GH32 enzyme, again validating our hypothesis that a requirement for growth during mono-colonization is the ability to break down polysaccharides that wild type EcN alone cannot degrade. Indeed, *in vitro* minimal media growth assays with a freshly transformed copy of this fragment enabled EcN consumption of sucrose [Figure 1f]. We further experimentally verified that of the fragments recovered in this experiment, this GH32-containing fragment uniquely enabled increased growth in media composed of the water-soluble extract of the Western diet, likely due to the high amount of sucrose (341 g/kg) in this formulation [Figure 1f].

To gain insight into the broad functional classes providing a selective advantage in each dietary condition, ORFs in each enriched fragment were classified into GO Pathways [Figure 2a]. In agreement with our qualitative observation of different fragments enriched in each condition, we also observed strong clustering of fragment function based on prebiotic supplementation. In all conditions, reflecting the “generalist” fragment that we recovered, “Carbohydrate Metabolic Process” was abundant (>23% relative abundance in all samples). In the presence of dextrin, we observed a unique additional signal for “Pyrimidine Nucleotide Biosynthetic Process”, reflecting a predicted glutamate synthase that reached high abundance. It has been previously demonstrated that *E. coli* has two primary pathways for glutamate synthesis, the glutamate synthase pathway and the glutamate dehydrogenase pathway, the latter of which is preferred during glucose-limited growth (Helling, 1998). As the dextrin monomer glucose is expected to be abundant in the dextrin condition, presumably due to the activity of the generalist GH32 enzyme, it was unsurprising that the glutamate synthase, and not the glutamate dehydrogenase pathway was enriched. In the presence of inulin, the major fragment (reaching 32-67% relative abundance in all mice at the end of the experiment) contained an ABC transporter that was classified as an “ATPase-coupled sulfate transmembrane transporter”. In the absence of prebiotic, the dominant fragment was the generalist glycosyl-hydrolase-containing fragment. Alone, no “specialist” fragment enabled increased *in vitro* growth on dextrin or inulin, or on Western diet extract broth. We conclude that in these EcN mono-colonized mice the dominant selective pressure remained the ability to enzymatically degrade long-chain polysaccharides, but that once this necessary function was fulfilled by a subpopulation of GH32-carrying EcN, as it would be by other taxa in a conventional gut community, secondary functions had room for enrichment in a host diet-specific manner.

**Figure 2.**
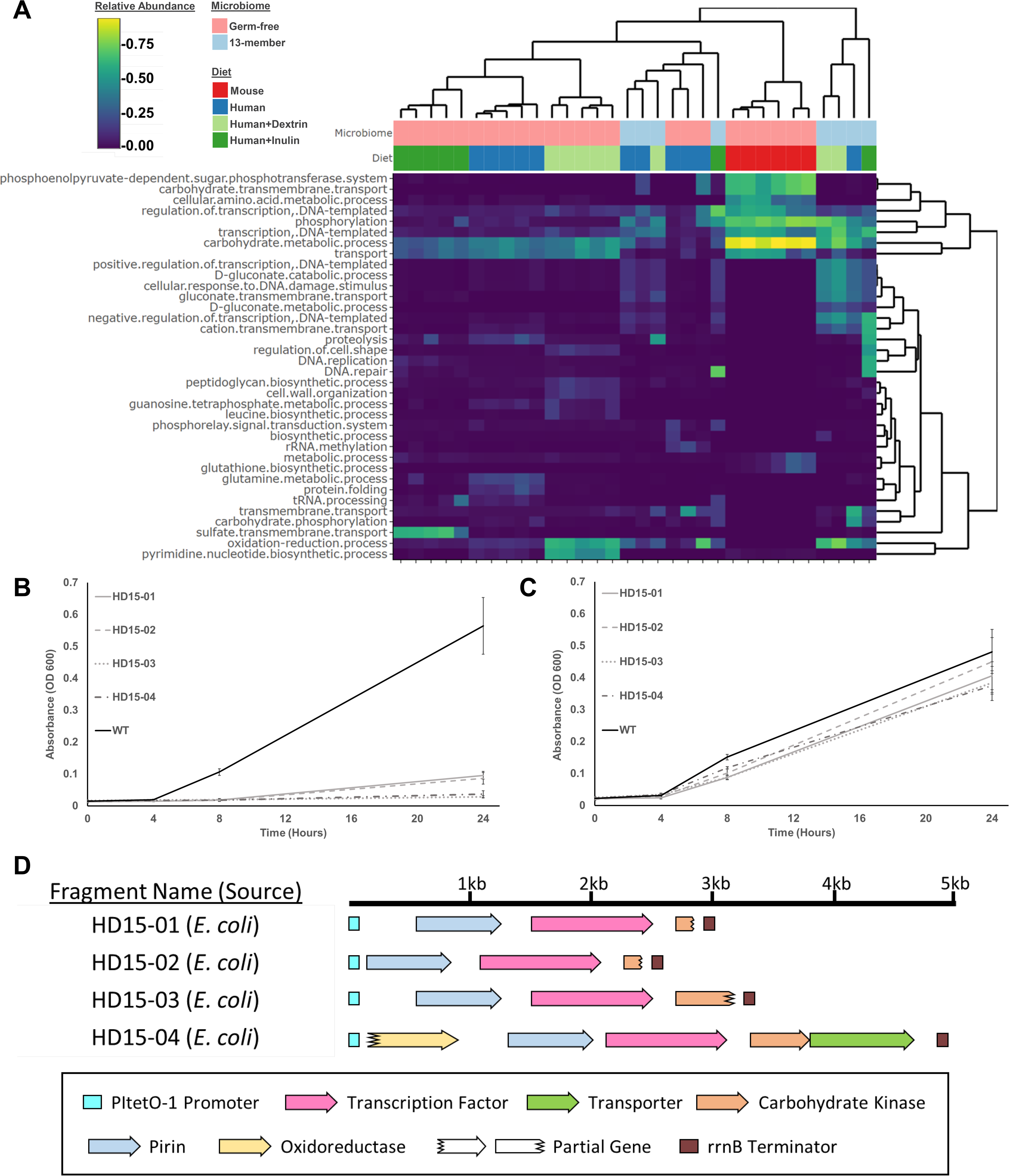
Comparison of functions enriched in functional selections in mice with different gut microbiome complexities. **A)** Heatmap of the relative abundances of metagenomic functions (GO classifications) enriched in EcN mono-colonized mice and mice pre-colonized with a 13-member defined community at the conclusion of the selection period. **B)** Growth curves of EcN strains harboring metagenomic inserts containing genes classified as a gluconate transmembrane transport or gluconate metabolic process (as depicted in A) on gluconate minimal media, or **(C)** glucose. **D)** Annotations of the metagenomic fragments characterized in B-C. Error bars represent the standard deviation of 6 replicates. See also Figure S1 and S3.

### Advantageous functions in a simplified microbial community

We then tested our hypotheses that i) in the presence of other taxa capable of degrading long-chain polysaccharides, this particular selective pressure on EcN would be relieved, and ii) with increasing complexity of the microbial gut community, EcN would be subject to selective pressures arising from direct competition and interactions with other microbes. Accordingly, 8 germ-free C57BL/6 mice were first colonized with a 13-member community of human gut-derived bacteria which broadly mimic the phylogenetic makeup of the human microbiome (Goodman et al., 2011), [Table S4]. After allowing this community to stabilize, the EcN metagenomic library was delivered via oral gavage and allowed to colonize for 5 weeks. Sequencing the metagenomic DNA contained in EcN after 5 weeks in the mouse gut revealed a highly diverse pool of fragments, relative to the mono-colonization experiments, supporting our hypothesis that in the presence of taxa expressing hydrolases that degrade complex polysaccharides, thus liberating di- and mono-saccharides for utilization by EcN, the selective pressure for acquisition of this family of hydrolases is relieved.

While no fragment reached a high abundance (>20% relative abundance) in all mice in the same experimental treatment, we did observe some functional convergence. The GO term “Carbohydrate Metabolic Process” was still consistently recovered across the majority of experimental replicates. One of the strongest signals we observed for condition-specific enrichment of a particular pathway was “Gluconate Metabolic Process”, with a transcriptional regulator, kinase, and transporter related to gluconate achieving >60% abundance in mice containing the 13-member defined community [Figure 2a]. We hypothesized that the presence of this pathway would directly alter the ability of EcN to metabolize gluconate, and tested this in *vivo*. When these fragments were cloned on a plasmid into wild type EcN, the recombinant strain was no longer able to grow on minimal media containing gluconate as the sole carbon source [Figure 2b], indicating constitutive activity of the gntR repressor present in these fragments. Gluconate utilization has been previously shown to be necessary for *E. coli* to colonize the mouse gut (Chang et al., 2004; Sweeney et al., 1996), so the consistent recovery of fragments reducing or abolishing gluconate utilization was puzzling. These past studies used conventional or streptomycin-treated mice hosting diverse mouse flora. In this study, it is possible that a different species in the 13-member human commensal consortium preferentially utilizes gluconate as a carbon source, making expression of the EcN gluconate utilization pathway metabolically wasteful, and providing a selective pressure for EcN to effectively downregulate the gluconate pathway by upregulating its repressor.

Together, these data indicate that the primary selective pressures imposed on EcN in the mammalian gut are related to carbohydrate utilization. Regardless of diet, there was a strong enrichment over the course of the functional metagenomic selections for glycosyl hydrolases that can degrade complex polysaccharides present in the diet, a function that in conventional gut microbiomes is provided by other taxa in the community. Once these taxa were reintroduced this selective pressure was relieved, but even in this case the most enriched functions were involved in carbohydrate utilization.

### Within-genome evolutionary change

To more thoroughly test our hypothesis that the main drivers of EcN adaptation in the mouse gut include factors involved in i) carbohydrate utilization ii) direct inhibitory interactions with other microbes, or iii) motility or adhesion in response to the mouse-specific mucins, we performed whole-genome sequencing on 401 isolates collected from mice that were either mono-colonized with EcN, colonized with a defined community, streptomycin treated, or conventional, and that in addition were either fed standard mouse chow, a Western diet, or a Western diet supplemented with dextrin [Tables S1-S2]. We also sequenced a reference EcN strain using both short and long read sequencing platforms, generating a high quality and manually curated closed assembly to which experimental isolates were compared. Among sequenced isolates, we observed a diverse array of genomic changes, with 456 total (336 unique) genomic changes distributed across 171 EcN genes [Figure 3]. The most highly-mutated strain had 5 SNPs, while the average strain had 0.86 SNPs/genome.

**Figure 3.**
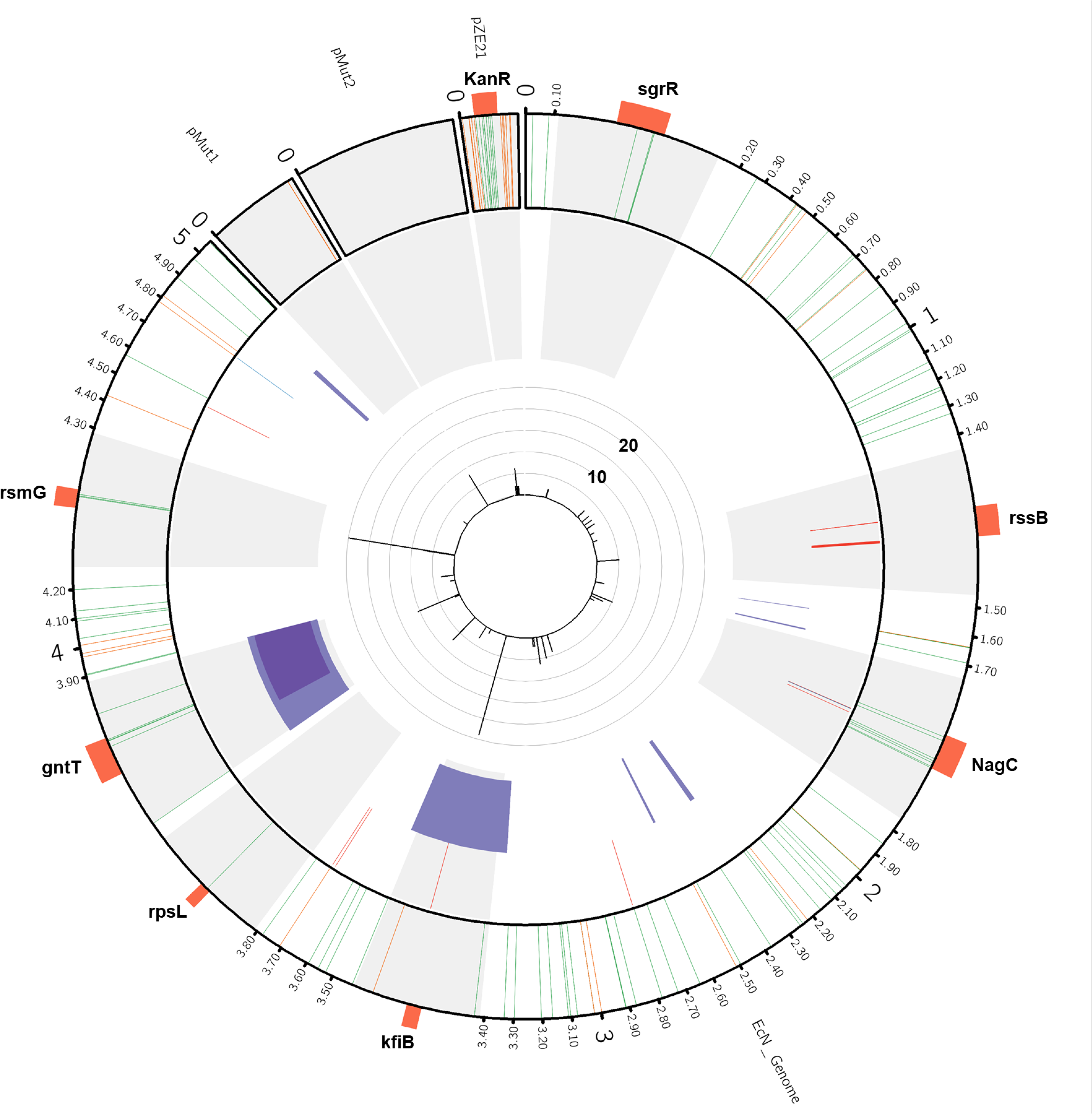
Summary of mutations detected in *in vivo* adapted EcN isolates. The EcN chromosome, its 2 native plasmids pMut1 and pMut2, and the expression vector pZE21 are depicted in a Circos plot. Light gray regions are on a scale 100x that of the rest of the chromosome for visibility. Genes of interest (orange) are labelled around the Circos plot. The outer track depicts all detected intergenic (orange) and nonsynonymous (green) SNPs. The next track depicts all small insertions (blue) and deletions (red). The third tack depicts all large deletions > 1kb (purple). Overlapping deletions are in dark purple. The inner track depicts the number of isolates that contained each mutation. Mutations found in only one isolate are excluded from this bar plot for visibility. Isolate counts for each mutation are capped at 25; this only affects the count for *rsmG* mutants, (167). See also Figure S4, Table S3, and Tables S5-S7.

### Adaptations involving nutrient utilization

Competition for nutrients plays a critical role in microbial community dynamics and colonization resistance (Pereira and Berry, 2017). Therefore, mutations that influence nutrient utilization in EcN are likely to give it a strong selective advantage, relative to wild type. In support of our hypothesis, we found that several genes involved in nutrient utilization were extensively mutated. The gene *gntT* was found to be mutated in 6 different ways across 21 strains isolated from four different mice, including 2 isolates with >10kb deletions containing this gene [Figure 4a]. The 4 other kinds of mutations included 3 unique early stop codons, and a D395N mutation. *GntT* encodes a high-affinity gluconate transporter (Porco et al., 1997). We hypothesized that *gntT* mutants isolated from these *in vivo* experiments would have an impaired ability to utilize gluconate as a carbon source, and so we tested the ability of representative isolates harboring these mutations to grow *in vitro* in gluconate minimal media. Isolates with the early stop codons were unable to grow, and the isolate with the D395N mutation exhibited an increased lag phase compared to wild type as indicated by the significantly lower optical density of its culture at 8 hours [Figure 4b]. When comparing all isolates from mice spanning the axis of microbiome complexity (mono-colonized to conventional), *gntT* mutants were significantly enriched in mice that had been pre-colonized with a 13-member defined community (p = 1.06E-6, hypergeometric test), although *gntT* mutants were also observed in EcN-monocolonized mice. Strikingly, this was the same phenotype recovered in the functional metagenomic selection in mice colonized with the defined community, where the strongest signal for functional enrichment was that of a gluconate utilization pathway whose phenotypic consequence was strongly inhibited growth on gluconate compared to wild type. This may be similarly due to preferential gluconate utilization by another member of the defined community balanced against the metabolic cost for EcN to maintain this pathway.

**Figure 4.**
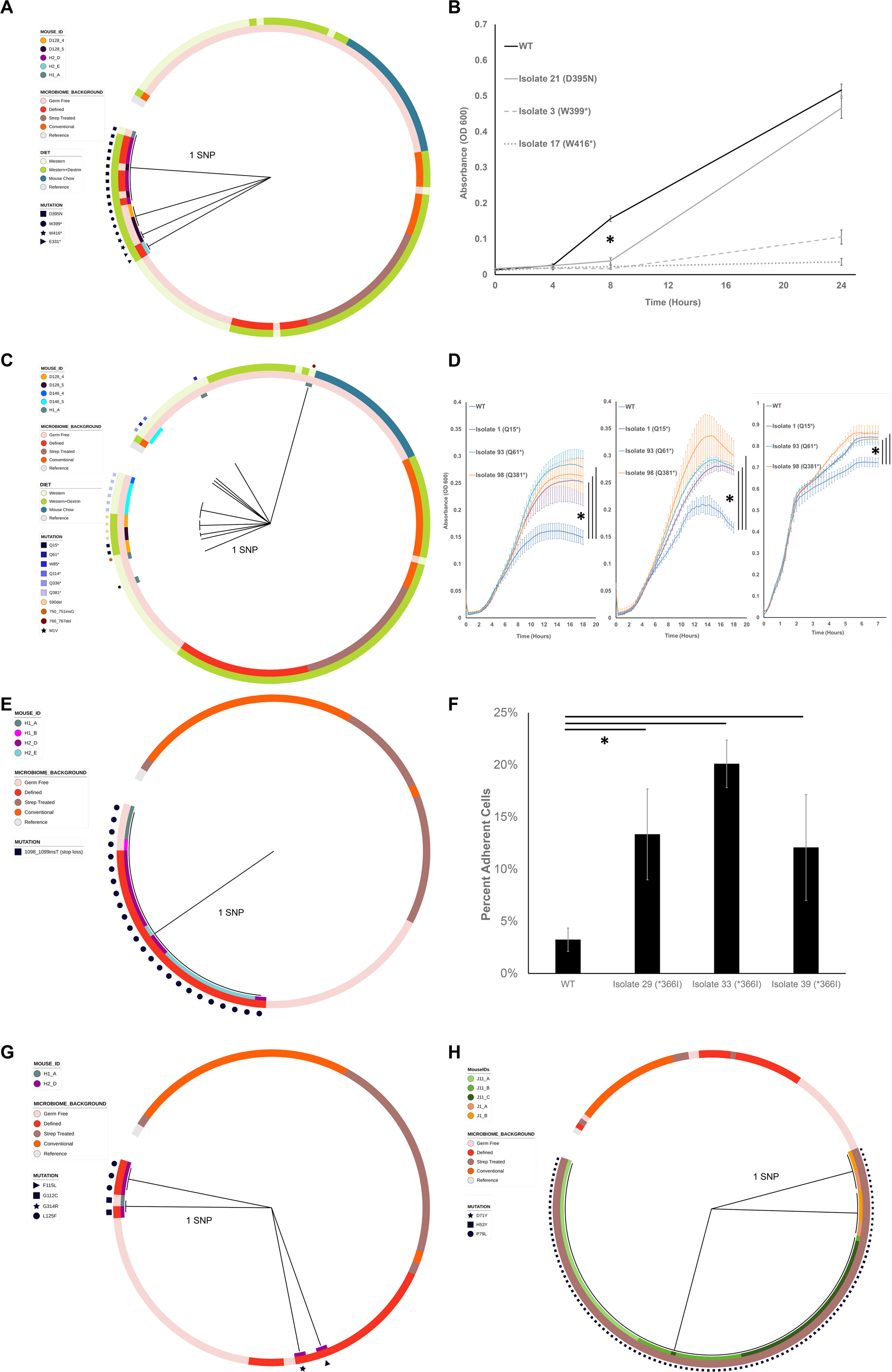
Phylogenies and phenotypic effects of mutant genes. Maximum parsimony phylogenies for adapted isolates based on their nucleotide sequences for **A)** *gntT*, **C)** *nagC*, **E)** *kfiB*, **G)** *sgrR*, or **H)** *rsmG*. Specific mutations are indicated. Color tracks depict the treatment conditions and the Mouse IDs from which isolates were isolated. The prefix of each Mouse ID indicates the cage in which the mouse was housed, e.g. Mouse H1_A and H1_B were housed together, but separately from Mouse H2_E. **B)** Growth curves of gntT mutants on gluconate minimal media. * P < 1×10-9, Student’s t-test comparing WT against Isolate 21 at 8 hours. **D)** Growth curves of *nagC* mutants on N-acetylglucosamine minimal media (left), Nacetylglucosamine and glucose minimal media (middle), and porcine gastric mucin and glucose minimal media (right). * P < 0.05, Welch’s t-tests comparing the area under the growth curves of each isolate and wild type, with Benjamini-Hochberg correction for multiple comparisons. Areas were determined by computing the integral of logistic models fit to the data for N-acetylglucosamine, and N-acetylglucosamine with glucose growth assays, or by numerically summing the area under the empirical growth curves for the gastric porcine mucin growth assay (growthcurvers R package). **F)** Percentages of *kfiB* mutant cells adherent to Caco-2 monolayers. Shown are the means of ratios of adherent CFU counts to total CFU counts (n = 3). Error bars are one standard deviation above and below the mean. * P < 0.05, Welch’s t-test with Benjamini-Hochberg correction for multiple comparisons.

Additionally, we found 10 different mutations in *nagC* in 19 isolates [Figure 4c]. These isolates were from 4 different mice used in the functional metagenomic selection for dietary conditions, and exclusively housed one of two metagenomic inserts, containing either GH32 or *gadX* transcriptional regulator, which may simply reflect the fact that those were the most abundant strains in the EcN population after the functional selection. All mutations resulted in either a premature stop codon, elimination of the start codon, or a frameshift. NagC is a repressor of N-acetylglucosamine and chitobiose import and utilization and is an activator of N-acetylglucosamine synthesis in its absence (Konopka, 2012; Plumbridge and Pellegrini, 2004). When mutated in this way, the pathways for N-acetylglucosamine and chitobiose utilization are likely permanently activated. We hypothesized that this may enhance the ability of EcN to consume host mucins, which have high levels of N-acetylglucosamine (Tailford et al., 2015). To test this hypothesis, we carried out an *in vitro* diauxic growth experiment and found that *nagC* mutants were able to switch more rapidly to growth on mucin under glucose limiting conditions than wild type EcN [Figure 4d]. We also found that *nagC* mutants were able to grow better on N-acetylglucosamine minimal media, and switch more rapidly to growth on N-acetylglucosamine under glucose limiting conditions [Figure 4d]. *NagC* mutants were exclusively isolated from germ-free mice that had been mono-colonized with EcN; such mutations enabling enhanced growth on a carbon source that is typically less preferred may be explained by the absence of taxa upon which EcN depends for polysaccharide degradation products. It has also been found that *nagC* is mutated during adaptation to osmotic stress (Winkler et al., 2014), which may provide EcN a selective advantage during transit through the GI tract.

### Adaptations involved in stress response

Probiotics are exposed to stressful conditions as integral parts of their life cycle. They must survive transit through the stomach and then take up residence in the nutrient-limiting and highly competitive environment of the GI tract. Our probiotics were subjected to selective pressure to remain viable in feces to gain repeated access to the GI tract through coprophagy. This is supported by the fact that stress response pathways were a strong target for natural selection in our experiment.

In particular, the gene *sgrR* was mutated 4 different ways in 8 isolates from 3 mice [Figure 4g]. These mutations were all nonsynonymous and did not result in stop codon formation. SgrR is a transcription factor which activates the expression of *sgrS*. SgrS is a small RNA which in turn inhibits the glucose-specific PTS transporter (Vanderpool and Gottesman, 2004). Under conditions of excess glucose-phosphate, SgrR is necessary to protect the cell from stress (Vanderpool and Gottesman, 2007). When comparing against all isolates from mice spanning the axis of biota complexity, *sgrR* mutations were significantly enriched in mice that had been pre-colonized with the 13-member defined community (p = 0.00732, hypergeometric test); however, because 6 of the isolates originated from one mouse that was in the defined community treatment group, this enrichment may be independent of the complexity of the microbiome. None of the mutations appear to be inactivating, and while a dN/dS analysis determined that there was not sufficient evidence to support positive selection, repeated nonsynonymous mutations in isolates from multiple, separately-caged mice suggests this stress response function is important for EcN fitness in the murine gut. Further, the *rssB* gene was mutated 4 different times in 9 different strains. Strikingly, all mutations to this gene were frameshift mutations. RssB regulates *rpoS* expression by directing it to the ClpX protease. RssB mutants display elevated RpoS levels (Battesti et al., 2011). RpoS is a master regulator of the stress response in *E. coli*, and *rpoS* mutants show enhanced resistance to stress (Fontaine et al., 2008).

### Adaptation in membrane-associated functions

The gene *kfiB* was mutated in 24 strains from 7 mice, each with the same loss of an early stop codon at amino acid 366, resulting in a full-length protein of 562 amino acids, indicating that in our wild type strain of EcN *kfiB* is either inactivated or truncated [Figure 4e]. When comparing isolates from mice spanning the axis of microbiome complexity, this potential gain-of-function variant was significantly enriched in mice that were pre-colonized with a defined community (p=3.01E-15, hypergeometric test). KfiB is a member of the K5 capsule polysaccharide (heparosan) biosynthesis cluster, which has been implicated in mediating the interaction between EcN and epithelial cells (Hafez et al., 2010; Leroux and Priem, 2016; Rigg et al., 1998). The role of *kfiB* in this gene cluster is not fully elucidated, but it has been shown to stabilize the *kfiAC* glycosyltransferase complex and be involved in heparosan export (Leroux and Priem, 2016). We hypothesized that our *kfiB* variants would exhibit altered adhesion to human epithelial cells compared to our wild type strain that harbors the early stop codon. To experimentally test this hypothesis, we carried out an *in vitro* adhesion assay with representative *kfiB* mutants and Caco-2 cell monolayers. As all *kfiB* variants had the same stop-loss mutation, 3 isolates were chosen as biological replicates. The results of this assay revealed that the loss of truncation in the kfiB mutants provided them significantly enhanced adhesion to Caco-2 cells [Figure 4f]. This loss of truncation may reflect re-adaptation of our reference EcN strain, which likely has been passaged often under laboratory conditions, to *in vivo* conditions and interactions with epithelial cells.

### Antibiotic resistance mutations

In colonization experiments in streptomycin-treated mice, we observed that the majority of isolates (167/204) contained mutations in *rsmG*, a 16S rRNA methyltransferase. *rsmG* is responsible for methylating G527 in the 530 loop of the 16S rRNA. The 530 loop is a target for streptomycin binding, and it has been shown that mutation of key residues in *rsmG* to alanine cause low-level resistance to streptomycin (Benitez-Paez et al., 2012). In particular, the three positions we identified with mutations (53, 71, and 79), were also mutated in prior work to give low-level streptomycin resistance and are predicted to be involved in catalysis, s-adenosyl-L-methionine (the methyl group donor) binding, and SAM binding, respectively. Unsurprisingly, when comparing isolates from mice that were treated with streptomycin to all other isolates, mutants were significantly enriched in mice that had been given streptomycin (p=1E-87, hypergeometric test). Analysis of mutations to this gene revealed strong evidence for extensive strain sharing among mice in the same cage. In particular, *rsmG* mutations clustered based on the cage from which the isolate was recovered. Isolates with the same mutation at position 53 were exclusively recovered from one cage (J11), while isolates with mutations at positions 71 and 79 were recovered from the other sampled cage (J1) [Figure 4h]. While the precise mechanism of microbiota transfer among humans is not known, this result indicates that if such a route is present, probiotics which evolved in one individual can be readily transferred to others. These results could indicate residual streptomycin present in the mouse gut, 1 day after delivery of the drug, and consequently that antibiotic treatment shortly before probiotic delivery may select for the presence of antibiotic-resistant probiotics.

We also observed 2 mutations to *rpsL* in 11 strains. All mutations occurred at residue 86 and converted it either to a cysteine or serine. These mutations only occurred in streptomycin-treated mice, and 10/11 times occurred in conjunction with a mutation in *rsmG*. Residue 86 is adjacent to residues that have been mutated previously to provide streptomycin resistance to EcN (Timms et al., 1992; Toivonen et al., 1999). From longitudinal sampling of 2 mice fed a Western diet and treated with streptomycin prior to EcN gavage, we calculate the mutation rate of EcN in the mouse gut to be 0.002 SNPs/genome/generation in the context of antibiotic treatment [Figure S4]. This rate is double that recently obtained for *E. coli* grown *in vitro* in the absence of antibiotic pressure (Lee et al., 2012). We expect that the combination of *in vivo* residence and antibiotic selection drove increased mutation rates in our experiment. Taken together, these findings are highly relevant considering the common practice of administering probiotics and fecal microbiota transplants after antibiotic administration (Hempel et al., 2012; Koenigsknecht and Young, 2013).

### *No* in vivo *plasmid acquisition*

We searched through our genome assemblies for contigs that did not map to the EcN genome, as this would indicate horizontal gene transfer of novel genetic material. Interestingly, we did not observe acquisition of new sequences over a 5-week timescale. The cause for this may be that EcN contains two endogenous plasmids (pMut1 and pMut2) (Grozdanov et al., 2004) in addition to the cloning vector that was used for this study. It is possible that these plasmids restrict incoming plasmids due to plasmid incompatibility groups or other defense mechanisms. In agreement with these results, *E. coli* Nissle has been shown to be a poor recipient of horizontally-transferred DNA, with a rate of DNA acceptance <10% the rate of *E. coli* K-12 for a variety of plasmid incompatibility groups (Sonnenborn and Schulze, 2009).

### In vivo *excision of large genomic sections*

In total, we observed 8 deletions larger than 1kb in size in 7 isolates. Two of these large deletions (24.3kb and 15.5kb in size) encompassed *gntT*, as further support that inactivation of this gene is beneficial for colonization in the context of a defined community. While EcN contains 5 putative phage regions (Arndt et al., 2016) [Table S5], only one (11.5kb) was observed to be excised *in vivo*. A 5.7kb deletion included a fimbriae locus. 3 deletions (23.3kb, 25kb, and 93kb in size) contained inverted repeat sequences, transposase, or integrase sequences, providing a potential mechanism for their excision. For example, the 25kb deletion, encompassing *glpR* (a ribokinase involved in ribose metabolism), *ydcZ* (an inner membrane exporter), *php* (a phosphotriesterase), along with various hypothetical proteins, was flanked by an insertion sequence 66 C-terminal element on one end and multiple transposases on the other. Interestingly, the 23.3kb and 15.5kb deletions were shared by the same strain. Copy number variation (CNV) analysis indicated that multiple isolates from the same mouse also exhibited copy number increases in two hypothetical proteins; a BLAST search revealed that these hypothetical proteins align significantly to an integrase and an inovirus Gp2 family protein. Together, these results indicate that large deletions or duplications occur in the EcN genome at low but significant rates during gut colonization, partially mediated by phage and transposon activity.

### Carbohydrate utilization, adhesion, and stress drive adaptations in EcN

Beyond the genes just described, we also performed GO annotation of all mutated genes. With 168 mutations, the GO pathway for Cell Division was the most commonly-mutated, but it should be noted that this GO pathway included the ribosomal genes mutated during streptomycin treatment. The next most abundant pathways were transcription (57 mutations), regulation of transcription (23 genes), and gluconate transmembrane transport (22 mutations) [Figure 5]. Collectively, this analysis revealed a diverse array of genomic changes undergone by probiotic EcN as it adapts to the mammalian gut. Our hypothesis that a main driver of probiotic adaptation in the gut is carbohydrate availability and utilization was supported; specifically, we have demonstrated that in the absence of other bacterial taxa the greatest selective pressure on EcN is glycosyl hydrolase activity. Ubiquitous mutations in kfiB provide support for our hypothesis that EcN will undergo adaptation specific to motility or adhesion in the mouse gut. Interestingly, other than one isolate with one mutation in its colicin receptor, we found no evidence of mutations involved in direct inhibitory interactions with other taxa such as toxins or secretion systems, but instead multiple signals for adaptation in stress response regulators. These data paint EcN as not an opportunistically offensive player, but instead a scavenger and survivalist.

**Figure 5.**
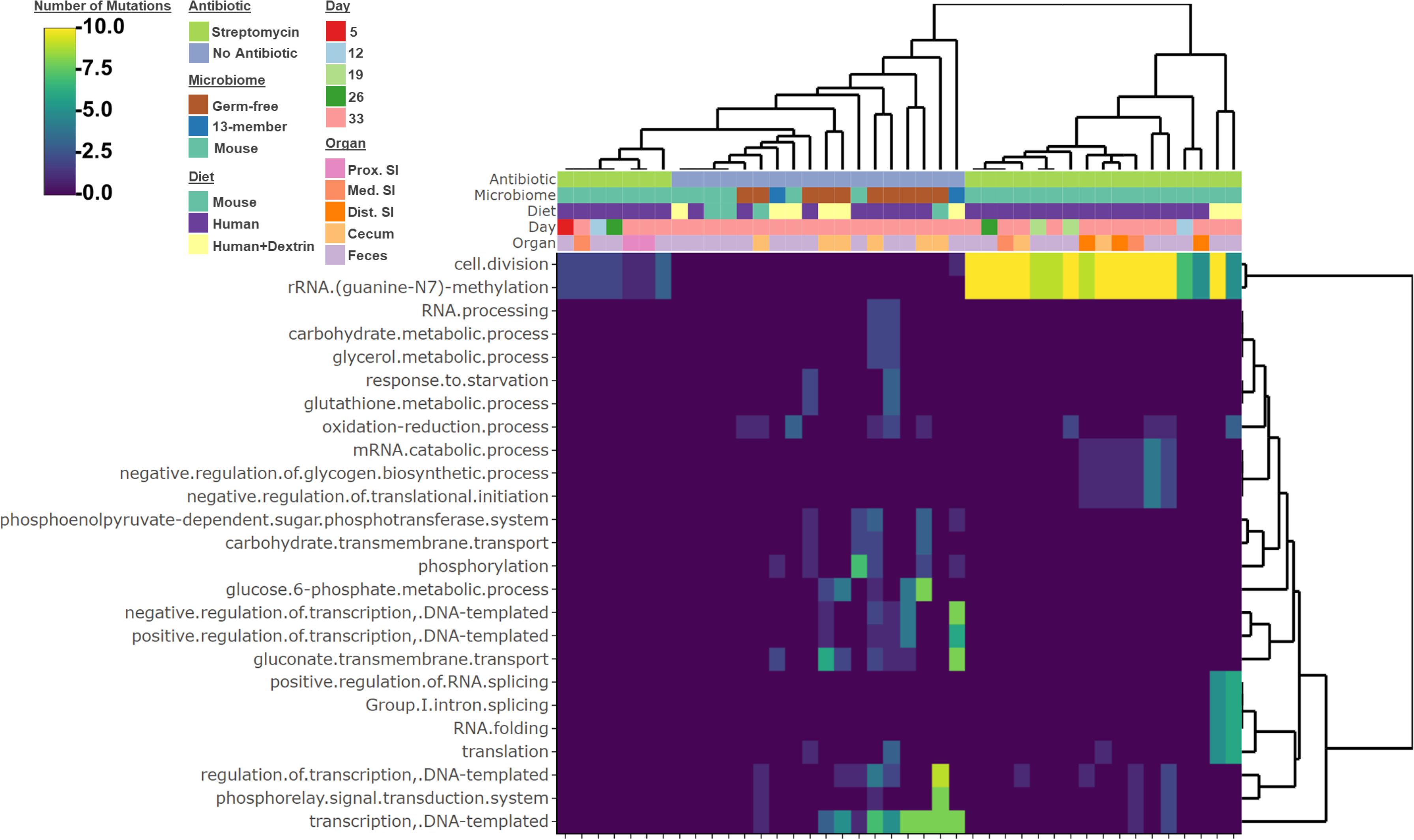
Comparison of functions mutated in EcN after passage through mice with a range of gut microbiome complexities and fed different diets. Heatmap of the cumulative number of nonsynonymous SNPs detected in genes grouped by their GO terms. Inclusion criteria for the heatmap was any function with at least two cumulative mutations identified across isolates from at least two mice. See also Figure S1.

### PKU treatment by engineered EcN

Engineered probiotics are gaining increased attention as low-cost delivery vehicles of drugs, microbial modulators, and metabolic functions (Hwang et al., 2017; Mansour and Abdelaziz, 2016; Palmer et al., 2018). Here we demonstrate the efficacy of EcN as a host for synthetic metabolism in the gut microbiota and determine whether mutations to engineered EcN accrue during treatment. To accomplish this, we turned to PKU, which is gaining increased attention as a metabolic disorder which may be treated by an engineered probiotic (Al-Hafid et al., 2015; Durrer et al., 2017; Falb et al., 2017).

PAH-deficient PKU creates a blockage in the tyrosine synthetic pathway leading to Phe accumulation in tissues. It has been shown that expression of a phenylalanine-metabolizing enzyme by probiotic *Lactobacillus* can lower the levels of circulating Phe by ∼33% in a mouse model of the disease (Durrer et al., 2017). We utilized PKU as a model to test the Phe catabolic properties of EcN. To this end, we expressed phenylalanine-ammonia lyase from *Arabidopsis thaliana* (McKenna and Nielsen, 2011) under the control of a weak, medium, or strong synthetic promoter (Anderson, 2006). The rationale for using different strength promoters is the known sexual dimorphism exhibited in the Pah^enu2^ mouse model, where females show moderately higher hyperphenylalaninemia than males (Sarkissian et al., 2008). We anticipate different levels of Phe reduction may be required to balance metabolic burden with equivalent therapeutic effect between the sexes. These expression cassettes were placed on a high copy plasmid and transformed into EcN. We observed strong rates of Phe consumption *in vitro* correlated to promoter strength, with the low strength promoter driving slower Phe consumption than the medium or high strength promoters [Figure 6a].

**Figure 6.**
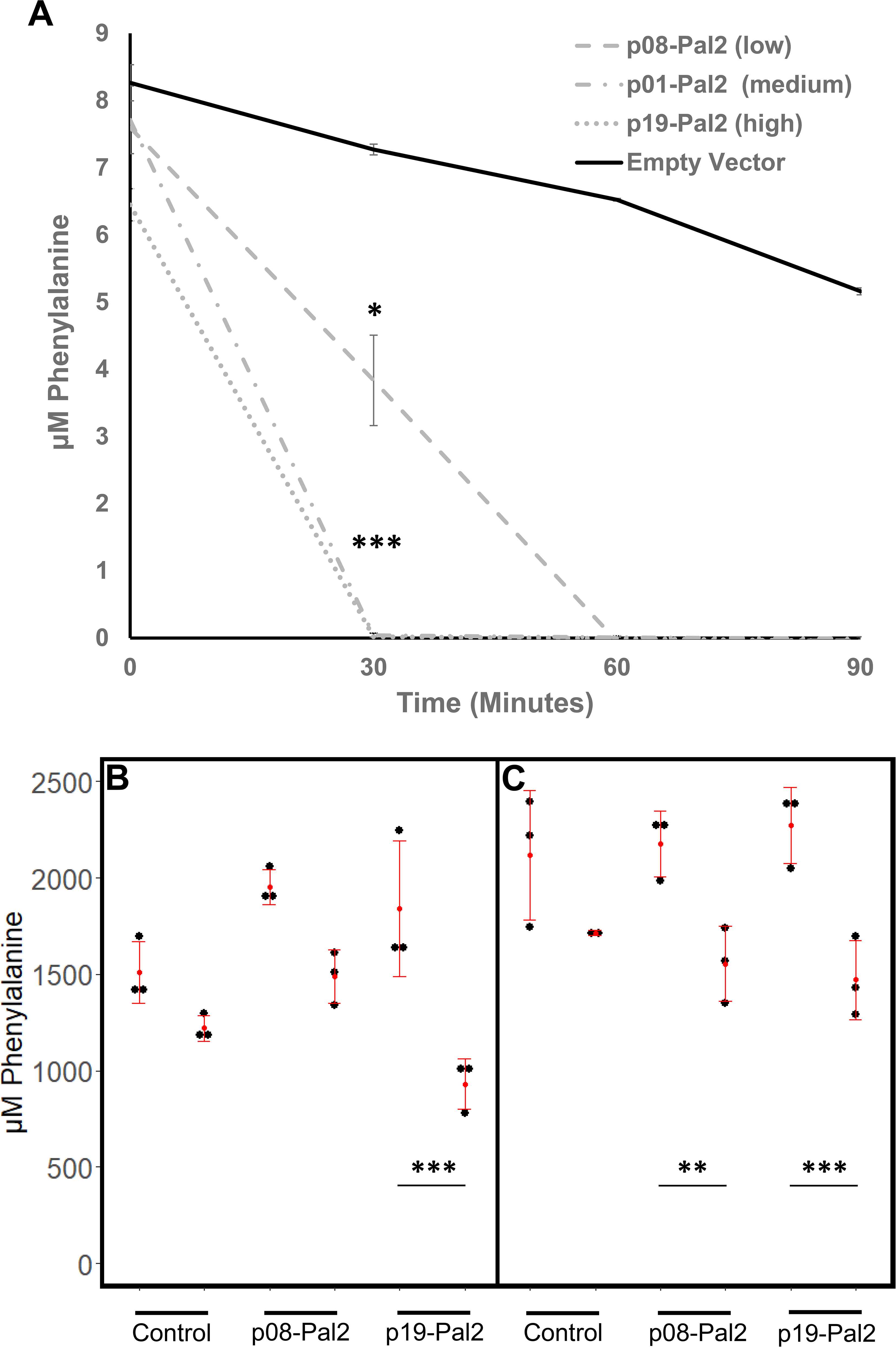
*In vitro* and *in vivo* degradation of phenylalanine by EcN:PAL2. **A)** *In vitro* phenylalanine concentration over time in minimal media supplemented with glucose and 10 μM Phe. Wells were inoculated with EcN expressing PAL2 under promoters of low, medium, or high strength. Shown are the mean of three technical replicates. Error bars represent one standard deviation above and below the mean. * P < 0.01, Welch’s t-test with Bonferroni correction for multiple comparisons. **B-C)** Paired serum phe concentrations for male (B) and female (C) homozygous mutant Pah^enu2^ mice before and 24 hours after gavage with wild type EcN or EcN expressing PAL2 under a high (p19) or low strength promoter (p08). Shown are means and standard deviations for three replicate mice. *** P < 0.0001, ** P < 0.01; P-values were calculated using the generalized linear hypothesis test with Bonferroni correction (glht function from the multcomp R package) on the coefficient estimates from a linear mixed-effects model generated by regressing Phe measurements on an interaction factor comprised of mouse gender, measurement timepoint (before or after treatment), and treatment (lme function from the nlme R package). Random intercepts by mouse were specified in the model.

To test whether the differences in promoter activity observed *in vitro* translate to activity differences *in vivo*, the strains with the high and low strength synthetic promoters were delivered to PKU mice via oral gavage (10^9^ cfu). Serum Phe was measured before and 24 hours after treatment. After 24 hours, male mice treated with EcN expressing PAL2 under a high strength promoter, and female mice treated with strains harboring either the high and low strength promoters, saw significant reductions in serum Phe concentrations [Figure 6b-c]. In male mice, serum Phe levels were reduced to half of their baseline levels 24 hours after probiotic delivery by the strongly-expressing probiotic [Figure 6b], indicating the potential of this strain for treatment of metabolic diseases. As expected, we observed dimorphism in the efficacy of this probiotic among males and female mice [Figure 6c]. We further observed *in vivo* differences in Phe reduction in keeping with the expected difference in activity of the low and high strength promoters, indicating the feasibility of tuning the performance of engineered probiotics *in situ*. We were able to recover EcN isolates containing the PAL gene one week after delivery, indicating that the recombinant plasmid was maintained by EcN in the absence of selection. Sequencing of recovered isolates failed to recover any mutations to the PAL2 gene over one week of treatment, indicating that this gene does not impose a substantial metabolic burden over this time course. We also only saw 1/16 strains accrue genomic changes. The two SNPs we observed in this strain were 1) a nonsynonymous mutation in the endoribonuclease ybeY and 2) a nonsense mutation in 4-alpha-glucanotransferase malQ. No other isolate sequenced during this study had mutations in these genes. Given the low frequency of the recovered mutations, we could not confirm our hypothesis that PAL2 expression confers a significant selective pressure to EcN in the PKU gut. Given the substantial amount of strain engineering that can readily be performed to further improve the efficacy of this treatment, we concluded that EcN represents a promising platform with which to treat PKU or deliver additional metabolic functions.

## Discussion

Unlike conventional abiotic therapeutics, the performance characteristics of probiotics can vary during treatment in response to natural selection. This variation can be relatively benign (i.e. increased residence time or loss of engineered function) or actively harmful (i.e. acquisition of virulence or antibiotic resistance), and therefore motivates detailed assessment. In this work, we described the evolutionary trajectories traversed by the candidate probiotic *E. coli* Nissle upon delivery to the GI tract.

We first simulated an upper bound for evolutionary rate through horizontal gene transfer via forced expression of a metagenomic DNA library derived from the healthy human fecal microbiota. By competing the resulting strains in the mouse gut, we found that the most advantageous fragments encoded nutrient utilization functions, rather than virulence factors, and that the specific nutrient utilization function was highly dependent upon host diet and microbiome composition. As probiotics are often administered to individuals with gastrointestinal or metabolic disorders, it would be interesting to determine whether these results extend to fragments sourced from dysbiotic microbiotas. While we recovered the highly conserved ABC transporters (Wilkens, 2015) in our experiments, our selections may have under-recovered other functions that require membrane translocation and display or secretion, due to unrecognizable signal sequences and incompatible translocation machinery. Alternate library construction strategies, such as massively parallelized fusion of robust translocation modules with metagenomic material, library expression enhancement with heterologous sigma functions, and expansion of heterologous host species, would be required to further explore this highly interesting functional class (Craig et al., 2010; Fleetwood et al., 2014; Gaida et al., 2015).

We next performed the most extensive analysis of *in vivo* probiotic genome evolution to date through whole-genome sequencing of 401 gut-adapted EcN isolates. We again observed strong selective signatures in nutrient utilization pathways, and even saw convergence between horizontal and vertical evolution in the inactivation of gluconate metabolism. In response to the harsh conditions of the gastrointestinal tract, we found that pathways involved in stress response were also significantly mutated. Surprisingly, we did not observe plasmid acquisition in any sequenced EcN isolate during gut passage, suggesting that the extent of HGT is low for this strain and validating the use of functional metagenomics to characterize in detail the heterologous functions conferring a selective advantage to EcN *in vivo*. In contrast to our expectation, functions involved in direct antagonistic interactions with other bacteria were not significantly mutated, collectively indicating that EcN defines its gut niche through metabolic competition. Finally, we did not observe any mutations in pathways associated with virulence, with the caveat that antibiotic exposure prior to probiotic delivery resulted in selection for antibiotic-resistant strains. Because probiotics and other commensals are often administered after antibiotic treatment to restore gut microbial diversity (Ciorba, 2012), refining the timescale over which antibiotics continue to exert selective pressure in the gut after administration is of critical importance.

Given the promising evolutionary characteristics of *E. coli* Nissle, we next demonstrated its potential as a delivery vehicle for added metabolic function, using phenylketonuria as a model disease. We found that expression of PAL was effective at reducing plasma Phe levels in male PKU mice by over 50% at 24h after administration, exceeding the reported efficacy of an engineered *Lactobacillus* (Durrer et al., 2017), and supporting engineered probiotics as a promising treatment route for PKU. This result also indicates that the powerful genetic tools available for *E. coli* Nissle manipulation position it well as a chassis for further engineering efforts, potentially enabling the generation of “personalized” probiotic therapies whose Pal expression is tuned to an individual’s unique Phe levels. Finally, we were unable to observe significant changes to the genome or plasmids of our engineered probiotic over one week of treatment, indicating that some types of engineered metabolism may not impose a substantial growth defect in the gut over the measured time scales.

Probiotics are an emerging class of therapeutics with increasing interest in diverse areas. Their use extends beyond the human large intestine and into habitats such as the mouth (Burton et al., 2006), skin (Wang et al., 2014), and urogenital tract (Stapleton et al., 2011). Microbes (natural and engineered) are also being deployed in agricultural (Berg, 2009), built environment (Liu et al., 2013), and natural (Pieper and Reineke, 2000) settings. Due to their wide use and propensity for genomic change we have just described, we argue that probiotics demand additional safety considerations beyond what is normally undertaken for traditional, nonliving entities. Coupled with continued development of biocontainment approaches (Chan et al., 2016), we envision the experiments described herein as a general pipeline to be applied to other probiotic strains in development to better understand their safety and engineering potential. It is possible to use *in vivo* functional metagenomic selections or genome evolution studies to develop engineered probiotics with increased colonization rates and longer residence times. However, the major selective pressures operating on any engineered probiotic may be substantially different than their wild type counterparts and would warrant further study of their *in vivo* evolutionary trajectories. As such, this work presents a generalizable framework for developing and regulating living therapeutics.

## Acknowledgments

We would like to acknowledge Chyi Hsieh and members of his lab for shared setup and maintenance of our germ-free mouse facility, Jessica Hoisington-Lopez, Eric Martin, and Brian Koebbe for next-generation sequencing and high-throughput computing support at the Edison Family Center for Genome Sciences and Systems Biology at Washington University in St Louis School of Medicine, and Thaddeus Stappenbeck, Tae Seok Moon, and members of the Dantas lab for helpful discussions relating to work described herein, This work is supported in part by the NIH Director’s New Innovator Award (http://commonfund.nih.gov/newinnovator/), the National Institute of Diabetes and Digestive and Kidney Diseases (NIDDK: http://www.niddk.nih.gov/), the National Institute of General Medical Sciences (NIGMS: http://www.nigms.nih.gov/), the National Institute of Allergy and Infectious Diseases (NIAID: https://www.niaid.nih.gov/), and the Eunice Kennedy Shriver National Institute of Child Health & Human Development (https://www.nichd.nih.gov/) of the National Institutes of Health (NIH) under award numbers DP2DK098089, R01GM099538, R01AI123394, and R01HD092414 to G.D. This work was also supported in part by the Kenneth Rainin Foundation Innovation and Breakthrough Awards (13H5) to G.D. N.C. received support from the NIDDK Pediatric Gastroenterology Research Training Program of the NIH, under award number T32 DK077653 (Phillip I. Tarr, Principal Investigator). A.F. received support from the Chancellor’s Graduate Research Fellowship Program at Washington University in St. Louis. A.J.G. received support from a NIGMS training grant through award number T32 GM007067 (Jim Skeath, Principal Investigator) and the NIDDK Pediatric Gastroenterology Research Training Program through award number T32 DK077653 (Phillip I. Tarr, Principal Investigator). M.W.P. received support from a NIGMS training grant through award number T32 GM007067 (Jim Skeath, Principal Investigator). M.K.G. received support as a Mr. and Mrs. Spencer T. Olin Fellow at Washington University in St Louis and from the National Science Foundation (NSF) as a graduate research fellow under award number DGE-1143954. The content is solely the responsibility of the authors and does not necessarily represent the official views of the funding agencies.

## Author Contributions

N.C., A.F., and A.J.G. designed and performed *in vivo* functional metagenomic selections in germ-free and conventional mice and carried out high-throughput sequencing of functional metagenomic hits. N.C. and A.F. also carried out high-throughput sequencing of adapted isolates, designed and performed *in vitro* microbiological experiments, designed and performed *in vivo* PKU experiments with Pah^enu2^ mice, analyzed data, and wrote the manuscript. M.P. and M.G. designed and performed *in vivo* functional metagenomic selections in germ-free mice. B.W. transformed metagenomic libraries into EcN and engineered a GFP-tagged EcN reference strain. S.S. carried out *in vivo* functional metagenomic selections in germ-free and conventional mice. Z.C. generated the experimental EcN:PAL2 strains for the PKU experiments. S.D. created and shared the Pah^enu2^ mouse model and provided guidance on its establishment and use. D.P carried out *in vivo* functional metagenomic selections in germ-free mice. G.D. designed experiments, analyzed data, and wrote the manuscript.

## Methods

### Media and Strains

*E. coli* Nissle was obtained as a kind gift from Dr. Phillip I. Tarr (Washington University in St. Louis School of Medicine). Unless otherwise indicated, EcN was grown in LB media under aerobic conditions at 37 °C. Agar was added at a concentration of 15 g/L for growth on solid media. For maintenance of the pZE21 plasmid (Lutz and Bujard, 1997) and its derivatives, kanamycin was added to a final concentration of 50 µg/mL after autoclaving.

Thirteen strains that were selected to mimic a healthy human microbiota composition were obtained as a kind gift from Dr. Andrew Goodman (Yale University) (Supplementary Table 4). Strains were cultured in TYG medium supplemented with D-(+)-Cellobiose (0.1% w/v; Sigma), D-(+)-Maltose (0.1% w/v; Sigma), D-(-)-Fructose (0.1% w/v; Sigma), Tween 80 (0.05% v/v; Sigma), Meat Extract (0.5% w/v; Sigma), ATCC Trace Mineral Supplement (1% v/v), ATCC Vitamin Supplement (1% v/v), N-butyric acid (4mM), propionic acid (8mM), isovaleric acid (1mM), and acetic acid (30mM) as in Goodman *et. al.* (Goodman et al., 2009). Culturing occurred under anaerobic conditions in a soft-sided plastic anaerobic chamber (Coy Laboratory Products). Stocks of strains were stored in E-Z crimp top vials (Wheaton) at −80˚C in Mega Medium with 20% glycerol. Stocks were titered by plating on BHI blood agar plates. Strains were pooled immediately prior to gavage such that each member was equally represented, and washed 3x in PBS. *Eubacterium rectale* and *Ruminococcus torques* displayed poor recovery rates from freezer stocks, so overnight cultures at early stationary phase were included in the final pool *in lieu* of stocks. Mice were gavaged with a total dose of 1×10^8^ CFU in a volume of 200 µL.

### Metagenomic library construction

Metagenomic libraries were sourced from a prior study examining the antibiotic resistome of the healthy maternal and infant microbiomes (Moore et al., 2015). Frozen glycerol stocks containing these libraries were thawed on ice, and plasmid DNA was extracted using a Qiagen Spin Miniprep kit. 100 ng of DNA was transformed into electrocompetent *E. coli* Nissle, and allowed to recover in SOC media for 1h at 37 °C. After 1h, 1 µL of SOC media was plated on LB+kan, to count transformants, and the remainder was placed in 50mL LB media containing kanamycin at room temperature overnight. After overnight growth, the library was centrifuged at 4000 rpm for 7 m, resuspended in 10mL LB+15% w/v glycerol, and frozen in 1mL aliquots at −80 °C. After freezing, one aliquot was thawed on ice, and serial dilutions were plated on LB+kan media to count viable cells. This procedure yielded a library containing 32.77 Gb of metagenomic DNA.

### Plasmids used in this study

All plasmids used in this study were constructed using a Golden Gate Assembly Mastermix from NEB, according to the manufacturer’s instructions. Plasmid pSPAL2At containing the PAL2 enzyme was obtained from Addgene (cat #78286), as a kind gift from David Nielsen (McKenna and Nielsen, 2011) Genbank files of all plasmids are provided in [Supplementary File 2].

### Mouse experiments

All mouse experiments were approved by the Washington University in Saint Louis School of Medicine Division of Comparative Medicine.

*E. coli* Nissle monocolonization experiments were performed in germ-free C57BL/6 mice (University of Michigan). Upon arrival, mice were provided food and water *ad libitum* for one week prior to colonization. For the mouse chow experiment, mice were provided autoclaved feed (Purina Conventional Mouse Diet (JL Rat/Mouse 6F Auto) #5K67) and autoclaved water. Western diet consisted of Envigo TD.88137, irradiated and vacuum-sealed. Inulin or Dextrin (Sigma) was provided in the drinking water at a concentration of 20 g/L, and autoclaved. 6 (for the mouse chow experiment) or 5 (for the Western Diet experiment) co-housed male mice were exposed to 10^8 colony forming units of *E. coli* Nissle via oral gavage of 100 µL cell suspension in phosphate buffered saline on day 1 of the experiment. Feces were collected at weekly intervals and immediately frozen at −80 °C. At the end of the experiment, mice were sacrificed through carbon dioxide asphyxiation and cecal contents were collected.

Experiments involving the 13-member synthetic microbiota were also performed in germ-free C57BL/6 mice (University of Pennsylvania Gnotobiotic Mouse Facility). Upon arrival, 3 germ-free mice (1-2 male, 1-2 female) per condition were housed separately in germ-free isolators. Mice were acclimated to their diet for one week prior to colonization with the synthetic microbiota. The synthetic microbiota was allowed to stabilize for one week prior to delivery of 10^8 colony forming units of *E. coli* Nissle via oral gavage of 100 µL cell suspension in phosphate buffered saline on day 1 of the experiment. Fecal samples were subsequently collected weekly for 5 weeks, and immediately frozen at −80 °C. At the end of the experiment, mice were sacrificed through carbon dioxide asphyxiation and cecal contents were collected.

Experiments involving the conventional mouse microbiota were performed in a specific-pathogen-free facility. 2 cages of 5 mice (Jackson Labs C57BL/6J) for each condition were maintained on the Western Diet with or without prebiotic for one week after arrival. After one week, mice were deprived of food and water for 4h, and given either 20mg streptomycin in 100 µL water, or 100 µL water alone via oral gavage. After this treatment, food and water was immediately returned. Daily over the next three days, 10^8^ CFU of EcN was delivered in 100 µL PBS via oral gavage. Fecal samples and intestinal contents were collected as above.

Experiments involving PKU mice were performed in a specific-pathogen-free mouse facility. For each condition (High PAL expression, Low PAL expression, or No PAL expression), 3 Male and 3 Female PKU mice (10-12 weeks of age) were co-housed and maintained on Phe-containing mouse chow. 100 µL blood was sampled from the femoral artery one week before the experiment to serve as a baseline for each mouse. Then, 10^9^ CFU of the appropriate EcN strain was delivered via oral gavage. One day after gavage, 100 µL blood was again sampled from the femoral artery. Fecal samples were collected 1 week after gavage. After collection, all blood samples were allowed to coagulate then centrifuged in BD Microtainer ® SST ™ tubes for 2 minutes at 10,000g to collect serum. Serum samples were then passed through 10 kDA molecular weight cut-off filters by centrifugation. Phe levels of the filtrates were determined using a Phenylalanine Assay Kit (MAK005, Sigma-Aldrich).

### Functional metagenomic library sequencing

DNA was extracted from fecal samples and intestinal contents using the PowerSoil DNA extraction kit (MoBio/Qiagen). 10 ng of this DNA was used as a template for a multiplex PCR with primers 1-6 [Supplementary Table 6] at equimolar concentrations using Taq Reddymix (Fisher Scientific). PCR recipe per sample was as follows: 12.5 µL Taq Reddymix, 3 µL primer mix (10 µM total concentration), 10 ng template, nuclease free water to 25 µL. PCR protocol was as follows: 94 °C for 10 m, 94 °C for 45 s, 55 °C for 45 s, 72 °C for 5.5 m, go to step 2 24 times, 72 °C for 10 m, 4 °C forever. PCR products were purified using a Qiagen Spin PCR Purification Kit. 500 ng PCR products in 200 µL elution buffer (from purification kit) were then placed in a AB1900 half-skirted plate (Fisher Scientific) and sonicated using a Covaris E220 sonicator. The settings on the sonicator were as follows: Peak Incident Power: 140, Duty Cycle: 10%, Cycles per Burst: 200, Treatment Time: 600 s, Temp: 7 °C. Sonicated products were cleaned using a Qiagen MinElute PCR purification kit according to the manufacturer’s instructions, and eluted in 22 µL elution buffer. 20 µL sonicated products were end-repaired through addition of 2.5 µL DNA ligase buffer with 10 mM ATP (NEB), 1 µL 1 mM dNTP, 0.5 µL T4 Polymerase (NEB), 0.5 µL T4 PNK (NEB), and 0.5 µL Taq Polymerase (NEB), and incubating for 30 m at 25 °C, followed by 20 m at 75 °C. Then, 0.8 µL T4 DNA ligase (NEB) and 5 µL pre-annealed sequencing barcodes were added to the full end-repair reaction, followed by 40 m at 16 °C, and 10 m at 65 °C.

Sequencing barcodes were designed according to the template described in Primers 7-8 (Supplementary Table 6), and synthesized by IDT in 2 96-well plates to a concentration of 500 µM. Each barcode was designed to be at least 2 mismatches away from every other barcode. 2 µL of each primer pair were added to 96 µL of TES buffer (10mM Tris, 1mM EDTA, 50mM NaCl, pH 8.0), and this mixture was diluted 1:10 in TES buffer, to a final concentration of 1 µM. Adapters were annealed by incubating at 95 °C for 1 m, followed by slowly cooling (0.1 °C per second) to 4 °C. Annealed oligos were kept cold and transferred to a −20 °C freezer until use.

Next, ligation products were size-selected to 300-400bp in size through gel electrophoresis. Briefly, the total ligation volume was loaded on a 2% (w/v) agarose gel in 0.5x TBE buffer using thin combs. This gel was run at 120V for 2h, or until the loading dye front reached ∼2/3 of the way to the end of the gel. DNA fragments between 300 and 400bp were excised, and purified using the Qiagen MinElute Gel Extraction Kit, eluting in 15 µL elution buffer.

Sequencing adaptors were then added to the size-selected ligation products through PCR. PCR recipe was as follows: 12.5 µL Phusion HF Mastermix (Finnzymes), 9.5 µL nuclease-free water, 1 µL 10 µM primer mix (Primers 9-10) and 2 µL gel-purified DNA. PCR conditions were as follows: 98 °C for 30 s, 98 °C for 10 s, 65 °C for 30 s, 72 °C for 30 s, return to step 2 17 times, 72 °C for 5 min, 4 °C forever. PCR products were then size-selected on a 2% agarose, 0.5x TBE gel as above. Sequencing libraries were quantified on a Qubit fluorimeter, and pooled at equimolar concentrations for sequencing. 100,000 2×150 paired-end sequencing reads were obtained per sample.

### Functional metagenomic sequencing analysis

Functional metagenomic contigs were assembled from sequencing reads and annotated using PARFuMS (Boolchandani et al., 2017; Forsberg et al., 2012). For each experiment, contigs were clustered at 95% nucleotide identity using CD-HIT (Fu et al., 2012). Then, vector-trimmed sequencing reads (from PARFuMS) were uniquely mapped to each contig using bowtie2 (Langmead and Salzberg, 2012), and mapping reads were counted using SAMtools (Li, 2011). Contig abundance was estimated as the fraction of reads mapping to each contig. To perform Gene Ontology mapping, open reading frames (from PARFuMS) were mapped against all nonredundant protein sequences using BLASTp (Boratyn et al., 2012), and annotated using InterProScan (Jones et al., 2014). Outputs of these two operations were then used as input to Blast2GO, according to manufacturer’s instructions. Each contig was associated with a GO annotation if at least one of its constituent open reading frames was. The abundance of each GO annotation was then estimated as the sum of contig abundances associated with that GO annotation. Heatmaps of GO annotations were plotted using the heatmaply package in R.

### Isolate sequencing

To isolate *in vivo*-adapted EcN isolates, intestinal contents or fecal samples were vortexed in phosphate-buffered saline and streaked on LB plates containing kanamycin. Single colonies were placed in liquid media and grown at 37 °C overnight. Un-adapted EcN strains were also grown overnight. Total DNA was extracted from each culture using the DNeasy UltraClean 96 Microbial kit (10196-4, Qiagen), and prepared for whole-genome sequencing using Nextera Tagmentation (Baym et al., 2015). Briefly, gDNA was brought to a concentration of 0.5 ng/µL in 1 µL volume. To each sample, 1.25 µL TD buffer, 0.125 µL TDE1 enzyme, and 0.125 µL nuclease-free water was added, and incubated at 55 °C for 15 m. Then, 11.2 µL KAPA HiFi master mix was added to each tagmented sample. Indexed sequencing adaptors (5 µM each) were thawed, and 8.8 µL was added to each sample. Then, PCR was performed using the following protocol: 72 °C for 3 m, 98 °C for 5 m, 98 °C for 10 s, 63 °C for 30 s, 72 °C for 30s, return to step 2 13 times, 72 °C for 5 m, 4 °C forever. PCR reactions were purified using AMPure XP beads according to the manufacturer’s protocol, and eluted in 60 µL resuspension buffer. Sequencing libraries were quantified using the Quant-iT PicoGreen dsDNA Assay Kit (Fisher Scientific), and pooled at equimolar concentrations for sequencing. 2 million 2×150bp sequencing reads were obtained per sample.

### MinION sequencing

Genomic DNA from 30 mL of overnight culture of control EcN was extracted using a Genomic DNA Buffer Set (19060, Qiagen) with the Qiagen protocol for Gram-negative bacteria (Qiagen Genomic DNA Handbook 06/2015) and the Genomic-tip 500/G (10262, Qiagen). Purified genomic DNA was sheared to a target fragment size of 10 kilobases using the Covaris g-TUBE™. All following steps were carried out in Eppendorf DNA LoBind tubes. Sheared DNA was repaired using the NEBNext FFPE Repair Mix (M6630), and repaired DNA purified from the reaction using Agencourt AMPure XP beads. DNA was then end-repaired and dA-tailed using the NEBNext End repair / dA-tailing Module (E7546) and again purified with AMPure XP beads. DNA was then prepared for MinION sequencing using the MinION Ligation Sequencing Kit 1D (SQK-LSK108). Briefly, adapters were ligated to the DNA using the MinION Adapter Mix (AMX1D) and NEB Blunt/TA Ligase Master Mix (M0367). Then the DNA product was purified using AMPure XP beads and the Minion Adapter Bead Binding Buffer (ABB). 350 ng of DNA product was then sequenced on a MinION R9.4 flow cell for 48 hours using the Running Buffer with Fuel Mix (RBF) and the Library Loading Bead Kit (EXP-LLB001). Average read lengths were 7 kilobases and the genome was sequenced with an average coverage of 50 reads.

### Isolate sequencing data analysis

Illumina sequencing reads were quality-filtered and adaptor-trimmed using Trimmomatic (Bolger et al., 2014). Isolate genomes were assembled with Illumina data using SPAdes (Bankevich et al., 2012). The control EcN genome (which had both Illumina and MinION data) was assembled using SPAdes in hybrid mode. Illumina reads were mapped to the assembled genome using bowtie2 and MinION reads were mapped using LAST (Frith and Noe, 2014). Mapping results were converted to indexed, sorted bam files using SAMtools and visualized using IGV (Thorvaldsdottir et al., 2013). The control EcN genome was then corrected manually by inspecting read alignments for dips in coverage, and correcting these regions using MinION data as well as homology to a published EcN genome (Reister et al., 2014) using an iterative process. After each iteration, Illumina and MinION reads were re-mapped to the corrected genome, and read alignments were inspected to verify an improved assembly. Coding sequences were annotated using Prokka (Seemann, 2014). One region of spurious assembly was due to imperfect tandem repeats present in the UpaH gene; this region was resolved by PCR amplification from the genome followed by Sanger sequencing. The result of this finishing process became the control genome to which all *in vivo*-adapted isolates were compared.

### Mutation Calling

Single-nucleotide polymorphisms and indels were annotated using two complementary approaches. The first method was Breseq, using default parameters (Deatherage and Barrick, 2014). The second method was VarScan2, where SNPs and indels were called if supported by a minimum read coverage of 30 and 80% abundance in those reads (Koboldt et al., 2012). VarScan2 variant call format output was then annotated with ANNOVAR (Wang et al., 2010). Mutations called by each method were manually verified through visualization of Illumina read alignments using IGV. Anecdotally, both methods did recover some false positive mutation calls, primarily due to the presence of a homologous metagenomic DNA insert causing spurious read alignments to the EcN genome. Phage sequences were identified using the Phaster web server (Arndt et al., 2016). To look for potential plasmid sequences, assembled contigs for each isolate were aligned to 1) all nonredundant nucleotide sequences, and 2) the control EcN genome using BLASTn. Our criteria for plasmid sequences was the following: >500bp in length, <90% of bases mapping to the EcN genome, no homology to the functional metagenomic insert present in the strain (if any), and no obvious contamination from other species. No contig satisfied these criteria, and therefore we concluded that no acquired plasmid sequences were obviously present in our *in vivo* adapted strains.

### Annotation

To perform Gene Ontology mapping, mutated open reading frameswere mapped against all nonredundant protein sequences using BLASTp, and annotated using InterProScan. Outputs of these two operations were then used as input to Blast2GO, according to manufacturer’s instructions. The abundance of each GO annotation was then estimated as the number of open mutated open reading frames with that annotation among the strains within a certain condition. Heatmaps of GO annotations were plotted using the heatmaply package in R.

### Copy Number Variation

Copy number variation was assessed using the CNVkit pipeline (Talevich et al., 2016). Briefly, short reads for each isolate were mapped to an indexed reference assembly using bowtie2 with the --very-senstitive-local option to generate SAM files. SAM files were then converted to sorted BAM files using Samtools. Coverage was calculated in bins using the CNVkit coverage function and compared to the coverage of the reference strain using the CNVkit reference function with the --no-edge option. Biases were adjusted using the CNVkit fix and segment functions. CNV hits were called if the bias-adjusted log2 copy ratio was greater than 1.3 or less than −1.3.

### Growth assays on functional metagenomic hits

Three biological replicates of EcN containing the appropriate plasmid were grown overnight in LB media containing kanamycin at 37 °C under anaerobic conditions. The cell densities of these overnight cultures were measured, and EcN was transferred to M9 MOPS minimal media containing the carbon source of interest at a starting optical density of 0.01 in a 96-well plate. BIOLOG plates were prepared with M9-MOPS minimal media per the manufacturer’s recommendations. A Breathe-Easy Sealing Membrane (Sigma) was placed over the plate. This plate was placed in a plate reader set to maintain 37 °C and read optical density (absorbance at 600 nm) every minute for 72h.

### Growth assays on mutant isolates

GntT mutants were grown on 0.4% gluconate minimal media. Three representative GntT mutant isolates were grown overnight in LB media at 37 °C aerobically, along with wild type EcN. The cell densities of these overnight cultures were measured, and the strains were transferred to M9 minimal media supplemented with 0.4% gluconate at a starting optical density of 0.05 in a 96-well plate as before. This plate was set in a plate shaker at 37 °C. Optical density (absorbance at 600nm) was measured at 0, 4, 8, and 24 hours.

NagC mutants were grown in diauxic growth experiments. As before, overnight cultures were transferred to M9 supplemented with 0.2% glucose 0.75% porcine gastric mucin (Sigma). Alternatively, overnight cultures were transferred to M9 supplemented with either 0.4% N-acetylglucosamine, or 0.2% glucose and 0.2% N-acetylglucosamine. These plates were set in a plate reader at 37 °C. Optical density (absorbance at 600 nm) was measured for 18 hours at an interval of 20 minutes.

### Epithelial cell binding assay

Caco-2 cells were seeded overnight in a 24-well plate at 50,000 cells/cm^2^ in DMEM-HEPES media supplemented with Glutamax (10564-011, ThermoFisher) and 10% Fetal Bovine Serum. They were incubated at 37 °C, 5.0% CO_2_ and 90% relative humidity. Overnight cultures of representative *kfiB* mutants or wild type EcN were grown in LB at 37 °C. The next day, the wells with Caco-2 cells were inoculated with mutant or wild type EcN that had been washed and resuspended in PBS. The multiplicity of infection was 1:100. The human and bacterial cells were co-incubated 37 °C, 5.0% CO_2_ and 90% relative humidity for 3 hours. Then, media was aspirated from each well. Three replicate wells for each strain were gently washed 3x with PBS, while three replicate wells for each strain were left untreated. All wells were then treated with 100 µL of 1% Triton-X for 10 minutes at room temperature to lyse the human cells. Then, 900 µL LB was added to each well and the cells resuspended by pipetting. These cultures were diluted and plated on LB agar plates and grown overnight at 37 °C. The next day CFUs were enumerated, and the percentage of adherent cells was calculated as the mean ratio of CFUs/mL from the washed and unwashed wells for each strain.

### In vitro *phenylalanine degradation assay*

EcN strains expressing PAL2 in the pZE21 plasmid under promoters of low (p08), medium (p01), or high (p19) strength, or harboring only empty vector, were grown overnight in LB at 37 °C. The next day the optical densities (absorbance at 600nm) of these cultures were measured and the strains were transferred to M9 minimal media supplemented with 0.4% glucose and 20 µM L-phenylalanine with a starting optical density of 1.0. Fractions of these cultures were removed at 0, 30, 60, and 90 minutes and immediately centrifuged. Supernatants were passaged through 10 kDa molecular weight cut-off filters and the phenylalanine in the filtrates was measured using a Phenylalanine Assay Kit (MAK005, Sigma-Aldrich). Phenylalanine levels were reported as average absolute nanomoles of phenylalanine per well. Three technical replicates were carried out for each strain.

### Gene phylogeny construction and enrichment testing

We searched for genes with variants significantly enriched in specific diet conditions or in specific microbiome complexities, using the hypergeometric test for enrichment and Bonferroni correction for multiple hypothesis testing. The significant genes included *gntT*, *nagC*, *sgrR*, *rsmG*, and *kfiB*. Gene phylogenies were created for these genes by aligning the gene sequence for all isolates involved in the comparison in which the gene variants were found enriched in a condition (that is, all isolates along the axis of different diets, or the axis of microbiome complexity). Alignments were generated using Clustal Omega. Maximum parsimony phylogenies for each gene were generated using the Dnapars program included in PHYLIP 3.695 (Felsenstein, 2005), and visualized using iTOL (Letunic and Bork, 2016).

### E. coli phylogeny construction

Representative E. coli genomes were downloaded from NCBI. PROKKA was used to generate general feature format (gff) files for each *E. coli* genome, including our assembled reference EcN genome (Seemann, 2014). Roary was used to generate a core gene alignment, using these gff files and the corresponding genome sequences (Page et al., 2015). The strain phylogeny was inferred with RaxML using the GTRGAMMA model and 1000 bootstraps (Stamatakis, 2014). The best tree was visualized using iTOL.

### Analysis of selection

*SgrR* was analyzed for evidence of positive or negative selection. The previously generated gene alignment and phylogeny were provided as input to the aBSREL, MEME, and FUBAR modules included in the HyPhy application hosted by the Datamonkey webserver (Delport, 2010). Neither of the three modules reported evidence of selection.

